# Neural engagement with online educational videos predicts learning performance for individual students

**DOI:** 10.1101/253260

**Authors:** Samantha S. Cohen, Jens Madsen, Gad Touchan, Denise Robles, Stella F. A. Lima, Simon Henin, Lucas C. Parra

## Abstract

Online educational materials are largely disseminated through videos, and yet there is little understanding of how these videos engage students and fuel academic success. We hypothesized that components of the electroencephalogram (EEG), previously shown to reflect video engagement, would be predictive of academic performance in the context of educational videos. Two groups of subjects watched educational videos in either an intentional learning paradigm, in which they were aware of an upcoming test, or in an incidental learning paradigm, in which they were unaware that they would be tested. “Neural engagement” was quantified by the inter-subject correlation (ISC) of the EEG that was evoked by the videos. In both groups, students with higher neural engagement retained more information. Neural engagement also discriminated between attentive and inattentive video viewing. These results suggest that this EEG metric is a marker of the stimulus-related attentional mechanisms necessary to retain information. In the future, EEG may be used as a tool to design and assess online educational content.

## Introduction

Student engagement is critical to academic success, and yet surprisingly little is known about the neural underpinnings of this process that lead to true psychological investment (Newmann, Wehlage, & Lamborn, 1992). Engagement is typically assessed with surveys after learning has commenced (Robinson & Hullinger, 2008), but these kinds of questionnaires do not provide a direct assessment of engagement during the learning process, and it is not clear that they are a reliable metric of psychological investment (Trowler & Trowler, 2010). In contrast to the classroom environment, where a teacher might readily assess physical manifestations of disengagement, the problem is much more acute in the increasingly common online learning environment, where expository lectures are often presented with videos (Means, Toyama, Murphy, Bakia, & Jones, 2009).

Online engagement can be measured by the number views or clicks (Koller, Ng, Do, & Chen, 2013), participation in online discussion forums (Brinton et al., 2014; Kizilcec, Piech, & Schneider, 2013), or by the length of viewing time (Guo, Kim, & Rubin, 2014; Kim et al., 2014). However, these outcome measures do not necessarily correlate with academic success (Koller et al., 2013). Online courses have an alarmingly high attrition rate of around 90% (Breslow et al., 2013; Jordan, 2014), indicating that students engage differently in online learning environments then they do in the classroom (Robinson & Hullinger, 2008). Furthermore, there is a large amount of heterogeneity in behavioral engagement with online courses (Kizilcec et al., 2013) and there is little agreement about how this behavioral data relates to psychological investment with the learning materials (Veletsianos, Collier, & Schneider, 2015). To address the difficulty in measuring engagement in the online learning environment, we leverage a method for assessing attentional engagement from brain activity during the process of learning and tie it to knowledge acquisition.

The approach builds upon findings that the similarity of neural responses are critical for both memory and engagement (Cohen, Henin, & Parra, 2017; Cohen & Parra, 2016; Dmochowski et al., 2014; Hasson, Furman, Clark, Dudai, & Davachi, 2008; Xue et al., 2010). Neural consistency can be measured either within a subject responding to repeated presentations of the same stimulus, or between subjects by measuring the similarity of their neural responses (Hasson et al., 2008; Xue et al., 2010). Since educational videos and classroom lessons are rarely repeated, we measure the reliability of neural responses of individual subjects by comparing them to other students, as they respond to the same educational videos.

We hypothesize that the similarity of electroencephalographic (EEG) responses across subjects to educational videos will be a sensitive measure of knowledge acquisition. When each individual’s brain activity is more similar to the group, their neural responses are more likely being driven by the stimulus, rather than by unrelated thoughts. To assess this similarity of individual subjects brain activity, we measured the inter-subject correlation (ISC) of EEG responses. ISC quantifies how similar each individual is to their peers. ISC is predictive of episodic memory, it is a sensitive metric of attentional state, and it is predictive of engagement when explicitly quantified by the scarce resources devoted to a video (Cohen et al., 2017; Cohen & Parra, 2016; Ki, Kelly, & Parra, 2016). Additionally, neural synchrony may be indicative of engagement in real-world classroom environments (Dikker et al., 2017). These previous studies did not explore how this “neural engagement” translates into an increase in learning efficacy. We predict that ISC will be indicative of both attentional state and learning performance in the context of educational videos. This study extends previous results by relating ISC to both incidental and intentional learning during the presentation of videos typically used in online education.

## Results

Knowledge acquisition was assessed in two cohorts who watched five educational videos on topics related to physics and biology. The first cohort (“intentional”, N = 18) took a short four alternative forced-choice questionnaire (9-12 questions) both before and after each video. The pre-video test (pretest) assessed baseline knowledge on the topic addressed by the video, and the post-video test (posttest) assessed facts imparted during the video and was therefore an assessment of semantic memory (Squire, 2004). Performance was much better on the post-test (67.4 ± 6.6 %) than on the pre-test (52.1 ± 5.4 %, t(17) = 5.9, p = 2e-5). The second cohort (“incidental”, N=21, 66.4 ± 6.6 %) watched the videos without knowledge of the post-test and took the same post-test as the first cohort after viewing all of the videos. Performance on the post-test was indistinguishable between the intentional and incidental learning groups (t(37) = 0.3, p = 0.8). Both groups performed above baseline level (44.6 ± 9.3 %, assessed from subjects who did not see the videos, N = 26, intentional: t(42) = 28.6, p = 4e-29, incidental: t(45) = 30.6, p = 1e-31). Two-way repeated measures ANOVAs, conducted separately in the incidental and intentional cohorts, with factors of subjects and movies, revealed a main effect for both factors in both cohorts (for videos: intentional: F(4,68)=11.1, p = 6e-7, incidental: F(4,80)=8.4, p = 1e-5, and for subjects: intentional: F(17,68)=3.6, p = 8e-5, incidental: F(20,80)=3.4, p = 4e-5). We predicted that “neural engagement”, as quantified by the ISC of neural responses, would explain some of the variability in test performance across subjects.

While subjects in both incidental and intentional cohorts watched the videos, EEG responses were recorded to assess the ISC evoked by the videos. ISC measures how well each individual’s EEG activity correlates with the other members of their cohort (intentional or incidental). The level of ISC significantly differed between the intentional and incidental learning paradigms (intentional: 0.11 ± 0.007, incidental: 0.08 ± 0.005, t(37) = 3.7, p = 7e-4), indicating that awareness of the upcoming test affected engagement with the videos. To measure the effect of attentional state on ISC, following Ki et al. (2016), subjects in both cohorts watched each video twice: while they were either attending normally or while they were disattending. Subjects first watched all videos and answered all questions (attend condition). After a short break, subjects watched all the videos again while silently counting backwards from 1000 in steps of seven (disattend condition). ISC was strongly modulated by the attentional state, and arguably the engagement, of viewers in both cohorts (Figure 1, intentional: 0.05 ± 0.007, t(17) = 13.1, p = 3e-10, incidental: 0.03 ± 0.004, t(20) = 12.3, p = 8e-11). Furthermore, the ability of ISC to discriminate between the attend and disattend conditions was assessed by the area under the receiver operating characteristic curve (AUC), a standard measure of discrimination performance, revealed that AUC was nearly perfect, and highly significant for both cohorts (intentional: AUC = 0.97 ± 0.03, incidental: 0.89 ± 0.03, mean ± standard deviation across all five videos, AUC measures how well individuals can be categorized as attentive or as inattentive, all p’s = 0.001). The attentional state of viewers was likely modulated by several factors including the counting task, fatigue, and motivation (see Discussion).

**Figure 1:**
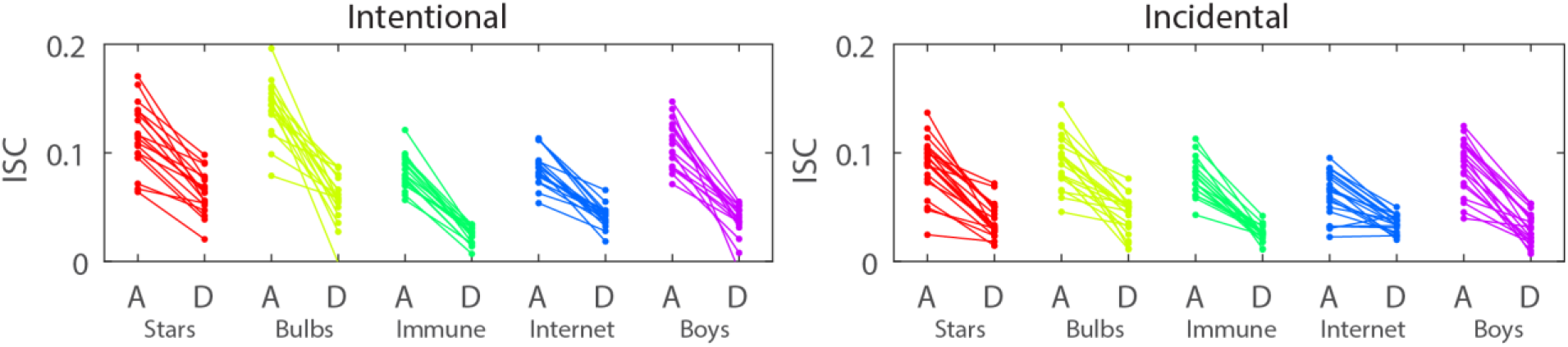
ISC is highly modulated by attention in response to educational videos. Inter-subject correlation (ISC) values for each subject (connected by a line) for each video while they either attending to (A) or were distracted from (D) the video’s content. ISC can discern attentional state in both the intentional and incidental conditions. Each video is portrayed in a different color (see Methods for video descriptions).

A three-way repeated measures ANOVA that compared ISC across attentional states, and repeated videos and subjects in each condition replicated the main effect for attentional state (intentional: *F*(1,176)=537.0, p = 5e-52, incidental: *F*(1,203)=468.4, p = 1e-51, Figure 1). The ANOVA revealed a main effect of video (intentional: *F*(4,176)=40.0, p = 6e-23, incidental: *F*(4,203)=10.0, p = 1e-7, Figure 1). Since ISC has been found to be a measure of neural engagement (Cohen et al., 2017; Dmochowski et al., 2014), this indicates that different movies interested subjects at different levels. There was also a main effect of subject (intentional: *F*(17,176)=4.8, p = 4e-8, incidental: *F*(20,203)=6.8, p = 1e-13, Figure 1), indicating significant inter-subject variability in engagement, just as there was for test performance.

To relate ISC, a neural proxy for engagement, to knowledge acquisition, ISC was used to predict performance on the post-video assessment. Figure 2A & B shows the relationship between ISC during the attentive state and test performance (Score [%]) for each subject for all five videos (indicated with different colors). When both ISC and test performance are averaged across the five videos, ISC correlated with test scores in both cohorts (Figure 2 C & D, intentional: r=0.57, p=0.01, N=18, incidental: r=0.41, p=0.07, N =21). Students with higher ISC, indicative of higher levels of neural engagement, also performed better on the tests querying their content. ISC in the disattend state did not significantly correlate with performance (intentional: r=-0.07, p=0.8, N =18, incidental: r=0.01, p=1, N =21). This drop in correlation suggests that ISC’s relationship with performance is contingent on attention, and does not easily result from other sources of variation across subjects.

**Figure 2:**
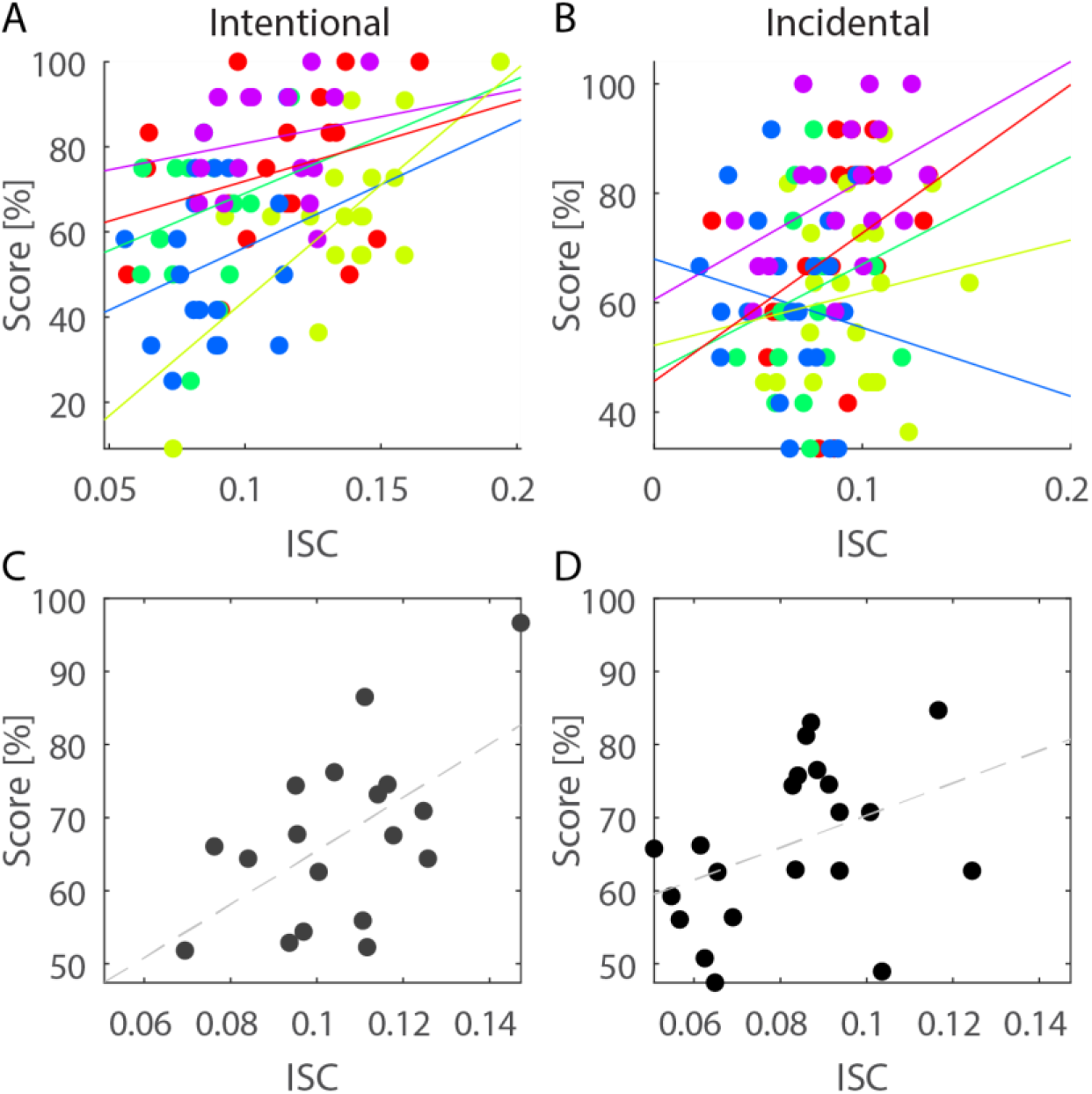
ISC predicts test performance for educational videos. A & B: Inter-subject correlation (ISC) and performance on the post-test (Score [%]) for all subject and all videos. Each point represents an individual’s ISC and test score for one of the five videos in either the intentional (A) or incidental (B) learning paradigm. Each video is indicated by a different color (colors are consistent with Figure 1). C & D: ISC correlated positively with test performance when both measures were averaged across all videos for each subject in both the intentional (C) and the incidental (D) condition.

## Discussion

The neural engagement evoked by educational videos was assessed with the inter-subject correlation (ISC) of the EEG recorded during video presentation. Subjects with high ISC elicit neural activity that is similar to their peers and they are therefore thought to be more engaged with the stimulus (Cohen et al., 2017; Dmochowski et al., 2014). ISC correlated with performance on a post-video assessment in both an intentional learning paradigm, in which subjects knew that they would be tested, and in an incidental learning paradigm, in which subjects were unaware of the test while watching the videos. After watching all videos and answering all questions, subjects watched the videos again while performing a distracting task. ISC could discern whether or not subjects were fully attending to the videos.

The high level of attentional modulation found with these educational videos is similar to levels found for entertaining narratives from conventional cinema (Ki et al., 2016). This result is surprising, because these putatively more “engaging” stimuli are theoretically more likely to capture attention. The evoked responses from these kinds of videos are therefore expected to be more consistent during an attentive state than during an inattentive state (Ki et al., 2016). The strong main effect of attention for both the incidental and intentional learning cohorts is likely the result of several factors. The disattend condition imposed a distracting task and it also consistently followed the attentive condition. Since the disattend condition always followed both the attentive condition and its associated tests, subjects were probably fatigued during this second presentation (in total the experiment lasted between 1 and 1.5 hours). This likely contributed to the nearly perfect discrimination performance of ISC. Additionally, in the intentional learning cohort, the attentive condition was book-ended by examinations, further motivating attention in this condition. Future studies should tease apart the task, fatigue, and motivation factors that induced the strong discrimination performance of ISC for the attentional state of viewers responding to educational videos.

ISC, a metric of attentional state, correlated with test performance in both the intentional and incidental learning groups. In both cohorts, the relationship between ISC and test performance was resolved with a smaller sample than that used previously ((Cohen & Parra, 2016) where effect size was r=0.49 compared to the values of r=0.57 and r=0.41 found here). This may indicate that ISC is a more sensitive metric of semantic knowledge acquisition, rather than the episodic memories that were previously tested (Cohen & Parra, 2016; Squire, 2004). It is also possible that since subjects were tested almost immediately after they watched the videos, their brain activity more strongly reflected their knowledge acquisition than in the previous experiment in which testing occurred three weeks after video exposure (Cohen & Parra, 2016). As students are often tested long after initial learning, future studies should consider testing subjects at longer latencies.

We designed the intentional and incidental learning conditions with the intention of modulating engagement with the videos. We expected that students would be more engaged when they expected an assessment. This effect was reflected by the modulation of neural engagement, but not by task performance, which was unaffected by awareness of the upcoming test. This curious dichotomy may indicate that the behavioral assessment was not robustly sensitive to the difference in attentional state, as indexed by the correlation of brain responses, across the two conditions. Although intentional learning is typically thought to result in better memories for words and pictures than incidental learning (Noldy, Stelmack, & Campbell, 1990), this is not the case when the items are processed more deeply (Schneider & Kintz, 1967). Although subjects were not instructed how to encode or process the material, the similarity in test performance between the two cohorts may result from a deeper level of processing inherently elicited by videos regardless of instructions (Craik & Lockhart, 1972).

The incidental learning condition is potentially more naturalistic than the intentional condition because students are typically not cognizant of an impending test when they are learning the material for it. However, even subjects in the intentional condition did not know which components of the video would be tested. Overall, the scenarios tested here are more realistic than previous efforts which have typically tested the memory for isolated words or pictures that are explicitly memorized (Atkinson & Shiffrin, 1968; Noldy et al., 1990). Future studies should extend these results beyond evaluations of factual information to more conceptual forms of learning. Although knowledge of facts may be the building blocks of more generalizable knowledge, testing this kind of learning may not correspond to the acquisition of more abstract understanding (Mayer, 2002).

In both intentional and incidental scenarios, neural engagement, as assessed by ISC, corresponded with memory strength. These results are consistent with the known link between attention and memory. ISC is therefore a potentially useful tool for relating video engagement to educational outcomes. The portable nature of EEG has obvious translational implications for this work in evaluating online learning materials and in real-world classrooms (Dikker et al., 2017; Poulsen, Kamronn, Dmochowski, Parra, & Hansen, 2017). Although some work has already been done that utilizes portable EEG in classroom settings (Dikker et al., 2017), none of this work has employed EEG as a measure of academic success. Here we have demonstrated a measure of EEG that is sensitive to both attentional state and knowledge acquisition. Since this research was done with educational videos, this finding is directly applicable to online courses where lectures are received by video. For massive open online courses (MOOCs), there is significant concern over low retention rates, which are attributed in part to lack of engagement (De Freitas, Morgan, & Gibson, 2015; Koller et al., 2013). ISC could be used to assess the engagement of learning materials in testing labs before they are disseminated. ISC has been shown to be predictive of the preferences of large populations (Dmochowski et al., 2014), it may therefore also be applicable to predict the educational efficacy of learning material for the broader population.

## Methods

### Stimuli

The five video stimuli were selected from the ‘Kurzgesagt – In a Nutshell’ and ‘minutephysics’ YouTube channels. They cover topics relating to physics, biology, and computer science that are not likely to be familiar to most subjects (Table 1, Range: 2.4 – 6.5 minutes, Average: 4.1 ± 2.0 minutes).

**Table 1:** Title, abbreviation, duration, web address, and example post-test question for the five videos used in experiment. *URL beginning with https://www.youtube.com/watch?v=

| Title | Abbreviation | Duration | URL-ending * | Topic area | Example Post- Test Question | Example post-test answer choices |
| --- | --- | --- | --- | --- | --- | --- |
| Why are Stars Star-Shaped? | Star | 3:28 | VVAKFJ8VVp4 | Physics | What is the shape of stars in Hubble space telescope photos? | 1. Four pointed<br>2. Concentric rings<br>3. Six pointed<br>4. Series of dashes |
| Why Do We Have More Boys Than Girls? | Boys | 2:48 | 3laYhG11ckA | Biology | Which of the following scenarios explains why the ratio of boys to girls changes with age? | 1. Boys are more likely to die from diseases at a young age<br>2. Girls are less likely to survive than males due to genetic reasons.<br>3. Girls more vulnerable to fatal diseases.<br>4. The sex ratio does not change |
| The Immune System Explained I – Bacteria Infection | Immune | 6:48 | zQGOcOUBi6s | Biology | What is the first cell type to intervene during a bacterial invasion? | 1. Guard cells / macrophage<br>2. Memory cells<br>3. B-cells<br>4. Dendritic cells |
| How Modern Light Bulbs Work | Bulbs | 2:57 | oCEKMEeZXug | Physics | The bulb does all of the following EXCEPT: | 1. Increases the brightness of the light emitted<br>2. Allows the filament to produce light more efficiently<br>3. Prevents gas from escaping from inside the bulb<br>4. Prevents oxygen from reaching the filament |
| Who Invented the Internet? And Why? | Internet | 6:32 | 21eFwbb48sE | Computer Science | Most internet traffic during the 1970's was: | 1. Written communication<br>2. Sharing digital images<br>3. Statistical data sets<br>4. Website retrievals |

### Experimental Design

Forty-two subjects participated in one of two experimental conditions: intentional or incidental learning. Twenty subjects participated in the intentional learning condition (six female, 23.9 ± 4.2 y.o., range 18 – 33, two subjects were eliminated because of poor signal quality). Twenty-two subjects participated in the incidental learning condition (seven female, 21.4 ± 1.2 y.o., range 20 – 25, one subject was eliminated because of poor signal quality). For both groups, videos were presented in a random order.

In the intentional learning condition, subjects were aware that they would be examined and they completed a short four alternative forced-choice questionnaire both before (pre-test) and after each video (post-test). The pre-test assessed background knowledge on the general topic covered in the video (9 – 11 questions). The post-test covered information imparted during the video (11 – 12 questions). Example post-test questions and answer choices are provided in Table 1 and a spreadsheet with all questions and answers is available at: https://tinyurl.com/y9xqu9bz. The incidental learning cohort completed the same post-test as the intentional learning group but after watching all five videos, and without prior knowledge of an assessment. After finishing all videos (attend condition) and corresponding assessments in the incidental and intentional cohorts, following a brief break (10 - 15 min), subjects watched the videos again (disattend condition), in the same random order as the attend condition. During this disattend condition, subjects were instructed to engage in the distracting task of counting backwards from 1,000 in decrements of seven (Ki, Kelly, & Parra, 2016). EEG was recorded during all quizzes and video presentations for both intentional and incidental cohorts.

To assess naive performance on the post-test, twenty-six subjects completed the post-test without seeing the videos (16 female, 22.5 ± 3.8 y.o., range 18 - 33). These subjects were drawn from the same experimental pool as the subjects who participated in the intentional and incidental conditions. All data collection and procedures were approved by the Institutional Review Board of the City University of New York.

### EEG Data Collection and Preprocessing

The EEG was recorded with a BioSemi Active Two system (BioSemi, Amsterdam, Netherlands) at a sampling frequency of 512 Hz. Subjects were fitted with a standard, 64-electrode cap following the international 10/10 system. The electrooculogram (EOG) was also recorded with six auxiliary electrodes (one located dorsally, ventrally, and laterally to each eye). All signal processing was performed offline in the MATLAB software (MathWorks, Natick, MA, USA).

Data pre-processing procedures followed Cohen & Parra (2016). The EEG and EOG data were first high-pass filtered (0.3 Hz cutoff), notch filtered at 60 Hz and then down-sampled to 128 Hz. After extracting the EEG/EOG segments corresponding to the duration of each stimulus, electrode channels with high variance were manually identified and replaced with zero valued samples. Eye-movement artifacts were removed by linearly regressing the EOG channels from the EEG channels, i.e. least-squares noise cancellation (Repovš, 2010). Outlier samples were identified in each channel (values exceeding 3 times the distance between the 25th and the 75th quartile of the median-centered signal) and samples 40ms before and after such outliers were replaced with zero valued samples. This zeroing procedure on a high-pass signal (zero mean) does not affect subsequent computations of correlation except for discounting correlation values by the fraction of zeroed samples.

### ISC and attention analysis

Inter-subject correlation (ISC) is calculated by first finding linear combinations of electrodes that are maximally correlated between subjects as they attend to the same video (Dmochowski, Sajda, Dias, & Parra, 2012). We refer to these combinations as correlated components, akin to principal or independent components. By construction, these linear combinations of electrodes are common to all subjects. The EEG data for each subject during each video is then projected into the space that maximizes correlation across subjects. This component extraction technique is a data-driven way to decide which of the electrodes are most informative about the correlation across subjects.

ISC is calculated for each subject as they watched each video by averaging the correlation coefficients of the projected EEG time-course between each subject and all other subjects. The ISC calculation implemented here is identical to previously published implementations and can be reproduced with code available at http://www.parralab.org/isc/ (Cohen & Parra, 2016; Petroni et al., 2018). Following previous research, ISC is calculated by using the sum of the three components of the EEG that capture maximally correlated responses between subjects (Dmochowski et al., 2012). For the attention analysis, the ISC components were trained from both the attend and dissattend conditions.

To test the difference between the attend and disattend conditions, student’s t-tests compared individual ISC values averaged across all stimuli for each subject. A three-way repeated measures ANOVA was used to compare ISC values for the repeated factors of movie and subject, and the nonrepeated factor of attentional state (attend vs. disattend). All statistical comparisons were computed within each cohort separately.

To test ISC’s ability to discriminate between the attend and disattend conditions, discrimination performance of ISC was assessed using the area under the receiver operating characteristic curve (AUC). The AUC analysis quantifies how well attentive and inattentive subjects can be distinguished by ISC. As such, each point on the receiver operating characteristic curve corresponds to one subject. Chance level of the AUC was determined by randomly shuffling the labels for the ISC values. Significance levels for each video and condition (incidental and intentional) were determined by comparing the AUC from the correct labels with 1000 renditions of randomized labels.

Test performance analysis and comparison with ISC

Test performance was assessed by calculating the percentage of correct responses on the pre- and posttests for each video. Student’s t-tests compared differences in the test performance between the pre- and post-tests and between the cohorts (incidental and intentional) after performance had been averaged across all tests for each subject. A two-way repeated measures ANOVA was performed with the repeated factors of movie and subject. ISC was compared to test performance by first averaging test performance across all post-video tests and averaging ISC values across all videos. A Pearson’s correlation value was then computed between ISC and test performance. Again, all analyses were conducted separately for each cohort.

## Acknowledgements

We would like to acknowledge SFN Grant number 603503991 for supporting this project.

